# Impact of Ebola virus nucleoprotein on VP40 virus-like particle production: a computational approach

**DOI:** 10.1101/2023.08.09.552607

**Authors:** Xiao Liu, Robert V. Stahelin, Elsje Pienaar

## Abstract

Ebola virus (EBOV) protein VP40 can assemble and bud in the form of virus-like particles (VLPs) when expressed in the absence of other EBOV proteins in mammalian cells. When nucleoprotein (NP) is co-expressed, VLPs will contain inclusion-body (IB) cores, and VLP production can be increased. However, the mechanism of how NP impacts VLP production remains unclear. Here, we use a computational approach to study NP-VP40 interactions. We find that NP may enhance VLP production through stabilizing VP40 filaments and accelerating the VLP budding step. Also, both the relative timing and amount of NP expression compared to VP40 are important for the effective production of IB-containing VLPs. We further find that there exists an optimum NP/VP40 expression ratio, and that earlier expression of NP compared to VP40 will produce IB-containing VLPs more efficiently. We conclude that disrupting the relative timing and amount of NP and VP40 expression could provide new avenues to treat EBOV infection. This work provides quantitative insights into how EBOV proteins interact and how those interactions could impact virion generation and drug efficacy.

## Introduction

Ebola virus (EBOV) is one of the most fatal known pathogens since its discovery in 1976 (1, 2). Over the last 40 years, more than 34,000 people have been infected and greater than 15,000 people have been killed in 44 known outbreaks (3, 4). While two antibody-based therapies were approved in 2021 (5, 6), the mortality rate is still greater than 30% even with therapy. There’s a need to continue developing new treatment for EBOV and better understand potential strategies for small molecule treatments.

Our understanding of the subcellular dynamics of EBOV is still limited. Research with live EBOV must be conducted in biosafety level 4 (BSL-4) labs, which slows research progress. Further, EBOV contains seven multi-functional viral proteins, all with complex protein-protein interactions, making it difficult to identify effective drug targets. Among them, matrix protein VP40 can assemble and bud in the form of virus-like particles (VLPs) from the plasma membrane of mammalian cells (7, 8). This feature of VP40 makes it a useful system to study the assembly and budding process of EBOV in BSL-2 conditions.

Aside from the viral matrix, the nucleocapsid (NC), which encapsulates viral RNA, is another important structure in EBOV assembly (9). The NC consists of at least NP, minor matrix protein (VP24), and polymerase cofactor (VP35) (10, 11). Apart from being wrapped by VP40, these NC-related proteins can modify the morphology and production efficiency of VP40 VLPs when co-expressed with VP40 (12, 13). To fully understand the assembly and budding process of EBOV, the complex interactions between matrix proteins and NC proteins must be considered.

NP, the critical component of the NC, has complex interactions with VP24 (10, 14), VP35 (10, 14), VP30 (15), VP40 (16, 17) and itself (11, 18, 19). It is also an important component of the ribonucleoprotein (RNP) complex, which is responsible for transcription, replication, and protection of EBOV RNA (20, 21). When expressed in mammalian cells, NP assembles into 20-25 nm diameter helical tubes, which have the same dimensions as the core structure of NCs (9, 11, 18). These findings suggest that studying NP assembly, in the context of VP40 assembly, can provide vital insights into the assembly of the NC and its interaction with the viral matrix.

Previous work has shown that co-expression of NP enhances VP40 VLP production (13). Though they found that the C-terminal domain (CTD) of NP is important for both increased VLP production as well as the recruitment of NP to VLPs, the detailed mechanisms of this influence of NP on VP40 VLPs remains unclear. Other studies found that cytoplasmic NCs contains no detectable VP40s when moving to the plasma membrane (22), and the movement speed of NC in the cytoplasm is not affected by VP40 co-expression (10). Besides, VP40 has the ability to recruit NCs to the site of budding on the cell membrane (10, 23). Together, these findings indicate that the interaction between VP40 and NP happens on the host cell plasma membrane. However, recent work suggests that the interaction between VP40 and NP takes place both in the cytoplasm and on the plasma membrane (two-stage interactions) (17). These authors concluded that NP interacts with VP40 in the cytoplasm through NP’s N-terminal domain (NTD) and induces a conformational change in NP’s CTD which is responsible for the recruitment of NP to the cell membrane and the incorporation of NP IBs into VLPs. When NP CTDs are mutated and lost the ability to interact with VP40s, they observed that VP40s will be trapped in IBs, as plasma membrane localization and VLP production will be reduced. These unknows and seemingly conflicting results demonstrate our lack of quantitative insights into NP-VP40 (or NP-VLP) interactions.

Mathematical modeling is a valuable tool to provide such quantitative insights into complex biological systems. We previously developed the first ODE-based model of VP40 assembly and budding at the intracellular level (24, 25). Our model suggested several mechanisms of VP40 and phosphatidylserine (PS) interactions regarding the formation VLPs, such as the influence of PS on VLP egress. We also revealed the dynamics of VP40 oligomers in the process of VLP assembly. This work illustrates the utility of mathematical models to generate hypotheses and guide experimental work.

Here, we build on our prior work with VP40 (cite both your papers if you can) to construct an ODE-based model of EBOV NP-VP40 interactions, assembly and budding at the subcellular level. We use this model to a) test which interactions between NP and VP40 can give rise to the experimentally observed impacts of co-expression, as well as b) quantify the impact of NP on VP40 VLP production.

## Results

### An ODE-based model replicates the impact of NP on VP40 VLP budding through a two-stage interaction

We developed the NP-VP40 model based on our previous VP40 model (24, 25). We incorporated the experimentally identified two-stage interaction between NP and VP40 by having cytoplasmic NP interacting with cytoplasmic VP40 dimer as well as full IBs interacting with membrane VP40 dimers (10). Our first aim is to see if our model can replicate experimental data from literature using this two-stage interaction assumption (13, 17).

Calibrated parameter sets successfully reproduced key experimental data. The addition of NP increases VLP production compared to VP40-only (Fig. 1A), while the membrane association-deficient NP mutant decreases both the membrane VP40 ratio and VLP production (Fig. 1B-C) compared to WT NP. Experimental work also indicated that the interaction between VP40 and NP in the cytoplasm should be increased when NP is mutated. In our model predictions, the increase (as indicated by P_1_, the portion of NP interacting with VP40 in cytoplasm) is consistent but very small due to the high P_1_ in WT NP group (Fig. S1, Table S4).

**Figure 1.**
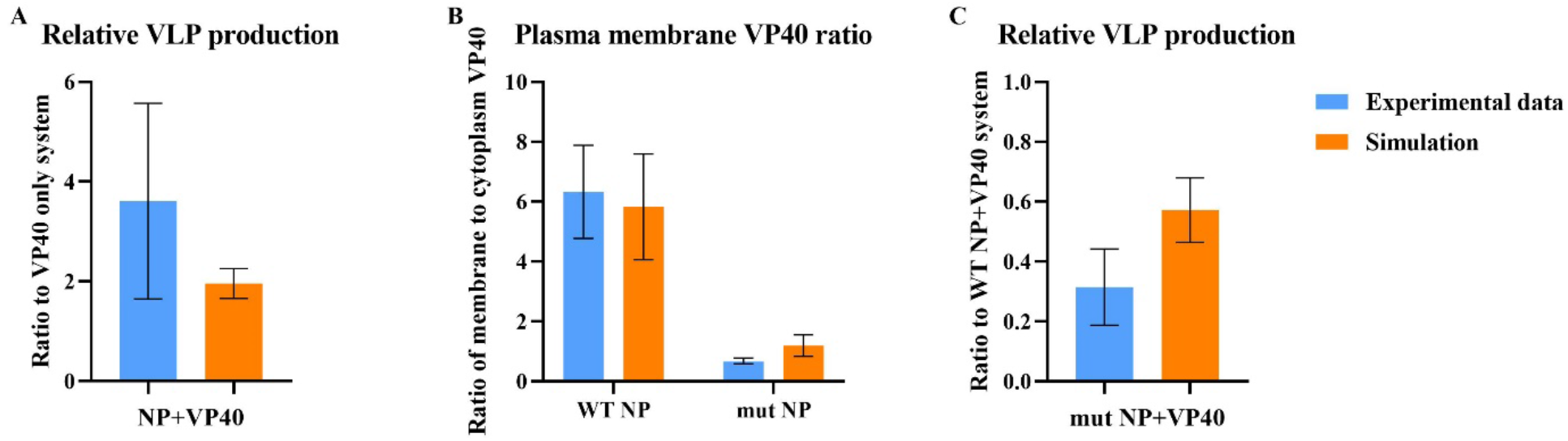
Simulation from NP-VP40 model. (A) Relative VLP production is increased to 196% on average when NP is co-expressed comparing to VP40-only. (B) Plasma membrane VP40 ratio is 5.83 when WT NP is co-expressed and reduces to 1.2 when mutant NP is co-expressed, on average. (C) Relative VLP production is reduced to 57.2% on average when mutant NP is co-expressed comparing to WT NP. Error bars indicate SD.

Finally, we evaluated our model against the bimodal size distribution of IBs when NP is expressed alone. Previous work also showed that the average size of IB will increase from 10-24 h, and both very small and large sized IBs are dominant, especially at later time points in a NP-only system (26). In their study, NP was detected after 10 h. However, it already becomes detectable in our model after 1h. Since there is no preparation stage for protein expression in our model, we believe there is a time shift between our system and their observation as the starting time of our simulation represents 9 h in their experiment. As a result, we looked at 1-15 h in our model instead. Our prediction agrees with the experimental observation. While the average size of IB is increased (Fig. S2A, Table S5), the distribution becomes binary after several hours (Fig. S2B, Table S6). Based on these quantitative and qualitative comparisons between our model results and experimental data, we believe that our model can replicate important features of the NP-VP40 system.

We used this calibrated and tested model to assess the impact of NP on VLP production, by looking at the VLP profile of the calibrated parameter sets. When NP is co-expressed with VP40, the increase in total VLP production compared to the VP40-only system (25) is attributed to the large amount of IB-containing VLPs. In fact, the IB-containing VLPs in the co-expression system outnumbers the total VLPs in the VP40-only system, while the IB-free VLP number is lower (Fig. 2, Table S7). This means co-expression of NP not only increases VLP production, but also prevents the formation of IB-free VLPs. We next wanted to study how NP influences the VP40 system to have such a VLP profile through local sensitivity analysis.

**Figure 2.**
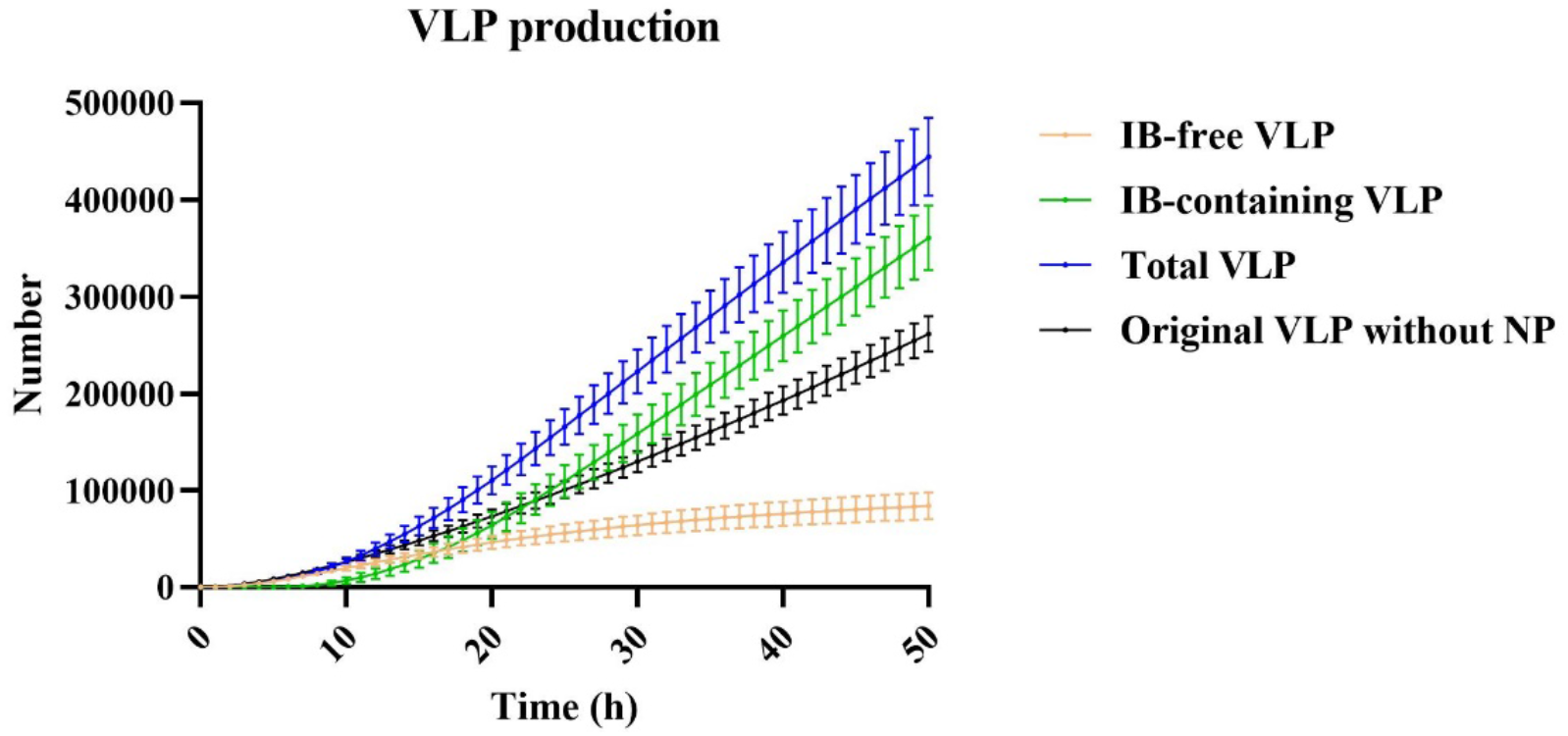
VLP production from NP-VP40 system. Total VLP production is much increased in NP-VP40 compared to VP40-only system, and the major form is IB-included VLP. On the other hand, IB-free VLP production is reduced compared to VP40-only system. Error bars indicate 95% CI.

### VP40 filament stability and VLP budding rate is positively impacted by NP

We next looked at how NP impacts the VLP filament assembly process. Our results indicate that the dissociation constants (k_D9,1_, k_D9,2_) for filament growth (nucleation and elongation, respectively) is decreased when NP is co-expressed with VP40 (Fig. 3, Table S8). This is expected, since VP40 can interact with IB, the stability of IB-bound filament is anticipated to be higher than IB-free filaments.

**Figure 3.**
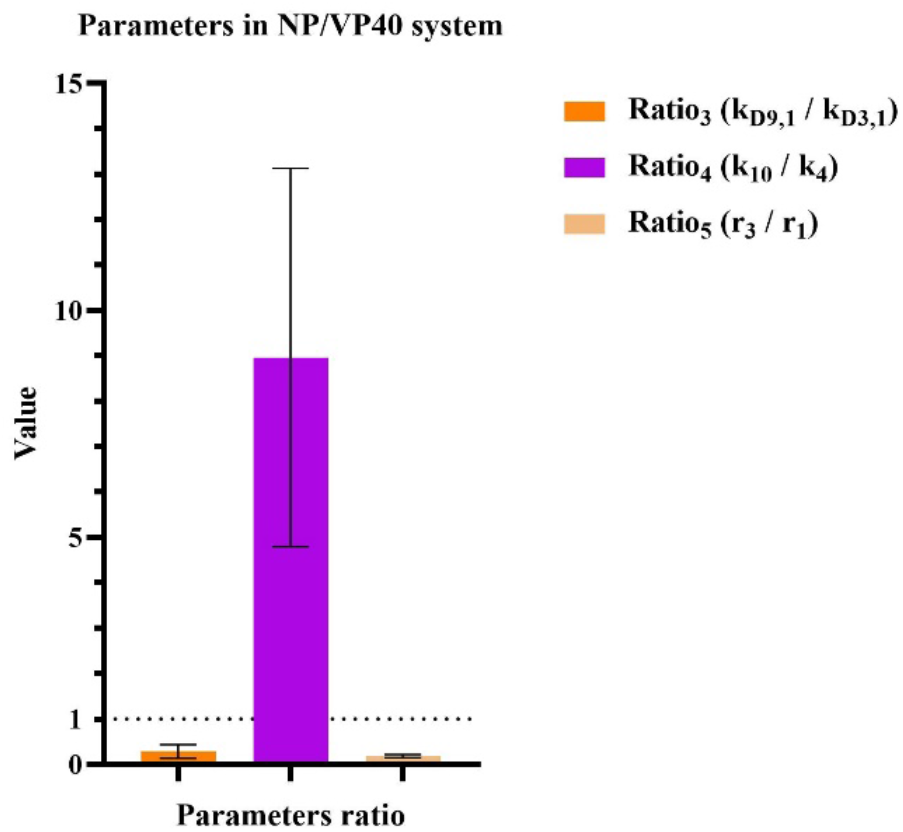
Range of important parameter ratios between NP+VP40 and VP40-only system. Co-expression of NP significantly decrease dissociation constant for filament growth and increase VLP budding rate. Monomer production rate for NP is much lower than VP40 in our system. Error bars indicate SD.

In order to study how the process contributes to the VLP profile, we then performed local sensitivity analysis for k_D9,1_ and k_D9,2_ (represented by k_9,1_’ and k_9,2_’). Results show that IB-free VLP production increases with k_D9,1_ and k_D9,2_ (Fig. 4A, Table S7, S9). In contrast, IB-containing VLP production is inhibited by both high and low k_D9,1_ and k_D9,2_ (Fig. 4B, Table S7, S9). This suggests that the impact of NP on increasing filament stability is critical for optimizing the proportion of IB-containing VLPs.

**Figure 4.**
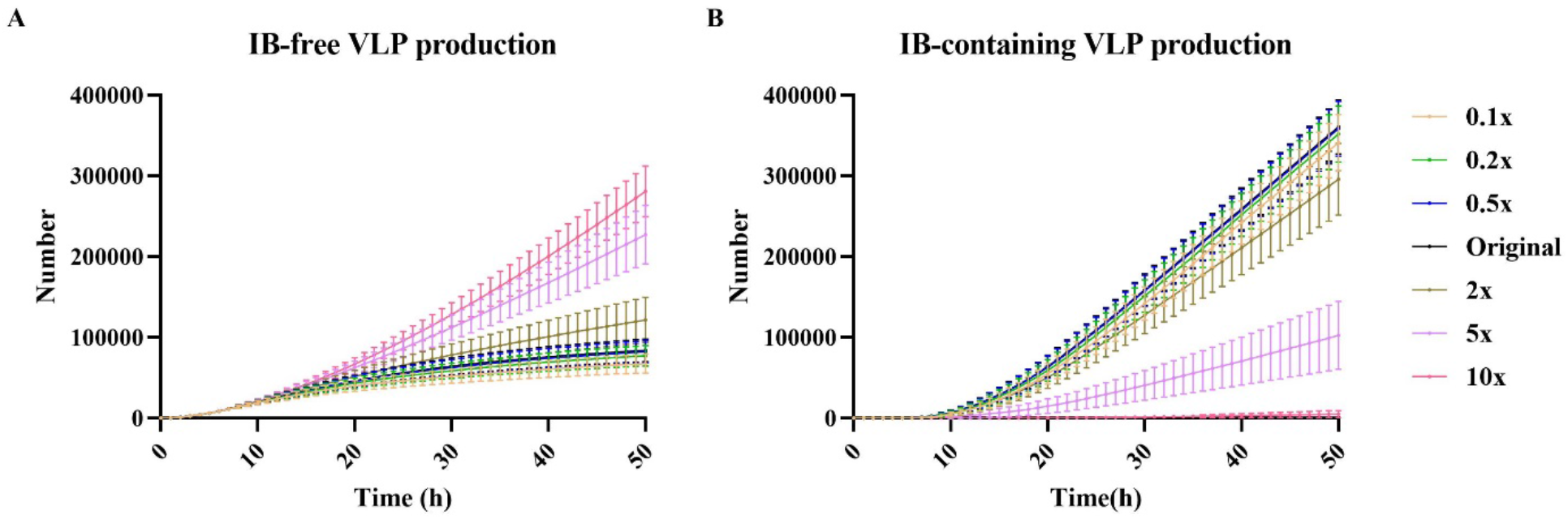
Influence of IB-containing filament growth dissociation constant on VLP production. (A) IB-free VLP production elevates as k_D9,1_ and k_D9,2_ increase from 0.1× to 10×. (B) IB-containing VLP production decreases when k_D9,1_ and k_D9,2_ are either very small or large. The reduction under large k_D9,1_ and k_D9,2_ is more obvious. Error bars indicate 95% CI.

We next evaluated how NP impacts the VLP budding process. Compared to the budding rate in the VP40-only system (k_4_), the VLP budding rate in the NP-VP40 system (k_10_) is increased (Fig. 3, Table S8). To study how IB-containing VLP budding rate contributes to the VLP profile, we performed local sensitivity analysis for k_10_. Unlike k_D9,1_ and k_D9,2_, the impact of k_10_ on IB-free VLP production is very limited (Fig. 5A, Table S7, S10). On the other hand, IB-containing VLP production increases with k_10_ (Fig. 5B, Table S7, S10). While the impact of IB on increasing VLP production is not important for IB-free VLP inhibition, it is critical for the promotion of IB-containing VLP production.

**Figure 5.**
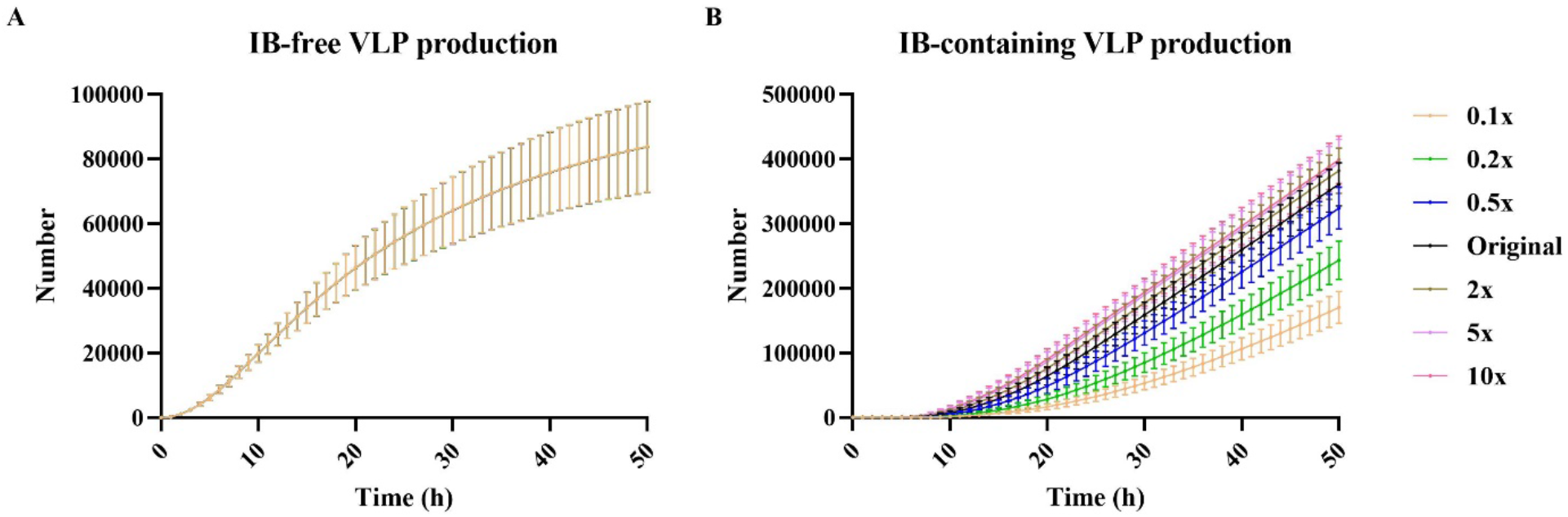
Influence of IB-containing VLP budding rate on VLP production. (A) IB-free VLP production is not influenced by k_10_. (B) IB-free VLP production elevates as k_10_ increases from 0.1× to 10×. Error bars indicate 95% CI.

Taken together, we conclude that NP will increase the stability of VP40 filament and VLP budding rate. The inhibition of IB-free VLP is related to the former impact, while the production of IB-containing VLP contains both influences. In terms of producing more effective virion particles, it’s important for EBOV to have as high as possible budding rate as well as an optimum stability for IB-containing filaments.

### NP/VP40 expression ratio and timing are both critical for the profile of VLP production

We also observed that the production rate of NP is lower than VP40 (Fig. 3, Table S8). To investigate how NP/VP40 production ratio impacts VLP production, we performed local sensitivity analysis for r_3_ (the NP monomer production rate constant). The production of IB-free VLP decreases as NP production rate increases, which is expected since NP can inhibit IB-free VLP production (Fig. 6A, Table S7, S11). However, IB-containing VLP production is also significantly inhibited when NP production is both too high and too low (Fig. 6B, Table S7, S11).

**Figure 6.**
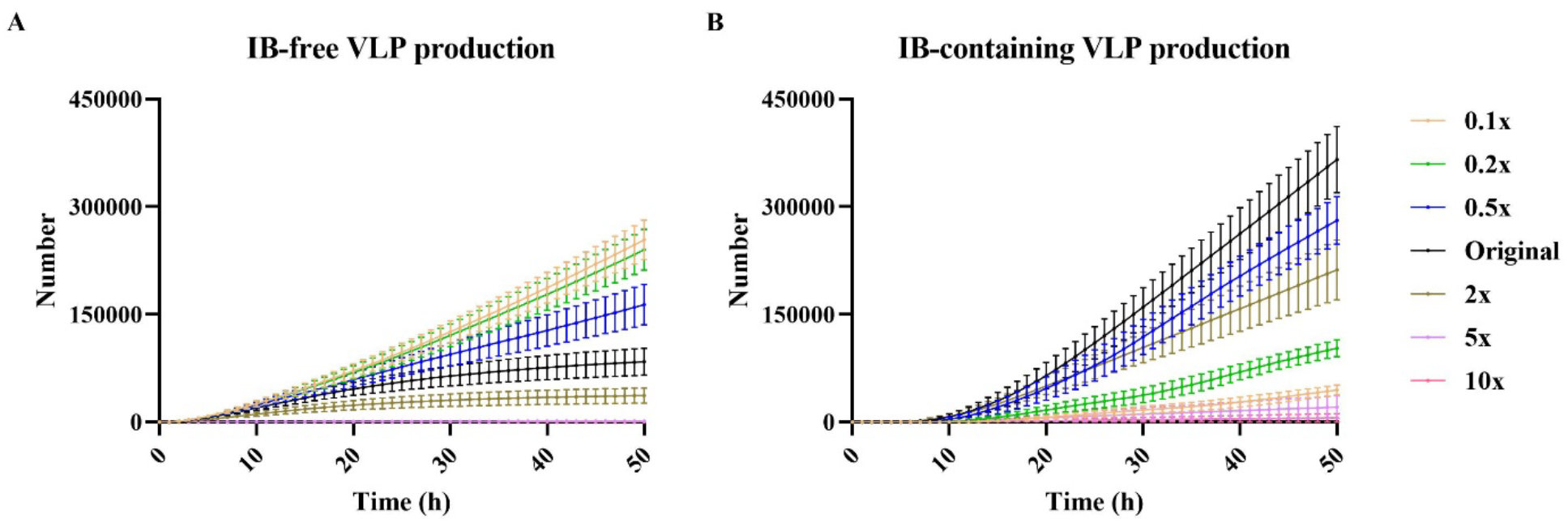
Influence of NP production rate on VLP production. (A) IB-free VLP production reduces as r_3_ increases from 0.1× to 10×. (B) IB-containing VLP production decreases when r_3_ is either very small or large. Error bars indicate 95% CI. 31 out of 50 groups are used for analysis as others have met tolerance problems in ode-solver.

We knew that too little NP will provide insufficient IBs for IB-containing VLPs and hypothesized that too much NP may trap large amounts of VP40 in the cytoplasm and prevent the formation of filaments, due to the two-stage interaction between NP and VP40. To test our hypothesis, we evaluated the VP40 bound to cytoplasmic IB. We do find that more VP40s will be trapped in cytoplasmic IBs as NP production rate increases (Fig. S3A, Table S12). However, when we looked at cell membrane VP40 number, though it is negatively related to NP production rate at the beginning, both low and high NP production rate will have more VP40 on the cell membrane at later time points (Fig. S3B, Table S12). This can be expected, since total VLP budded in those groups are low (Fig. S3C, Table S12), especially in high NP production groups (5×, 10× NP production rate), leaving more VP40s remaining on cell membrane.

The question remains why VLPs are not being budded when membrane VP40 concentration is so high. To answer this question, we further evaluated the concentration of filament building blocks (VP40 dimers). The cell membrane VP40 dimer concentration is very low in high NP production rate simulations (Fig. S3D, Table S12) while there are more IBs moving to cell membrane in those groups (Fig. S3E, Table S12). These IBs moving to the cell membrane are less likely to “release” VP40 dimer since IB-containing filaments are more stable than IB-free filaments (Fig. 3, Table S8). Thus, while the demand of cell membrane VP40 dimer is high for the maturation of those IB-containing filaments, the concentration of VP40 dimers is low, which leads to low IB-containing VLP production. Taken together, when NP production is too high, VP40 will be trapped in both cytoplasmic IB and incomplete IB-containing filaments that are unable to bud. Note that in this model, we assume all NPs in the cytoplasm have the ability to interact with VP40. If this is not true, the optimum NP/VP40 expression rate for higher IB-containing VLP production can be much higher.

Apart from the expression ratio between NP and VP40, we also wanted to determine the impact of the expression timing of NP relative to VP40 on the VLP profile. To do this, we performed the expression time test for the two proteins (see Methods). This analysis demonstrates that the later NP is expressed relative to VP40, the more IB-free VLP will be produced (Fig. 7A, Table S13). IB-containing VLP production reaches a peak when NP and VP40 are co-expressed at the same time, followed closely by the NP-5h earlier expression group (Fig. 7B, Table S13). We then calculated the energy efficiency of VLP production by dividing the IB-containing VLP production by total protein production. Co-expression of NP and VP40 at the same time has the highest efficiency initially (since no IB-containing VLP can be produced at the time for other groups), then gets surpassed by the NP-5h earlier expression group (Fig. 8, Table S14). Considering both IB-containing VLP production amount and energy usage efficiency, we believe that it is most beneficial for EBOV when VP40 is expressed a bit later than NP. This is aligned with the genome sequence of EBOV, as NP is closer to the 3’-end compared to VP40 (27).

**Figure 7.**
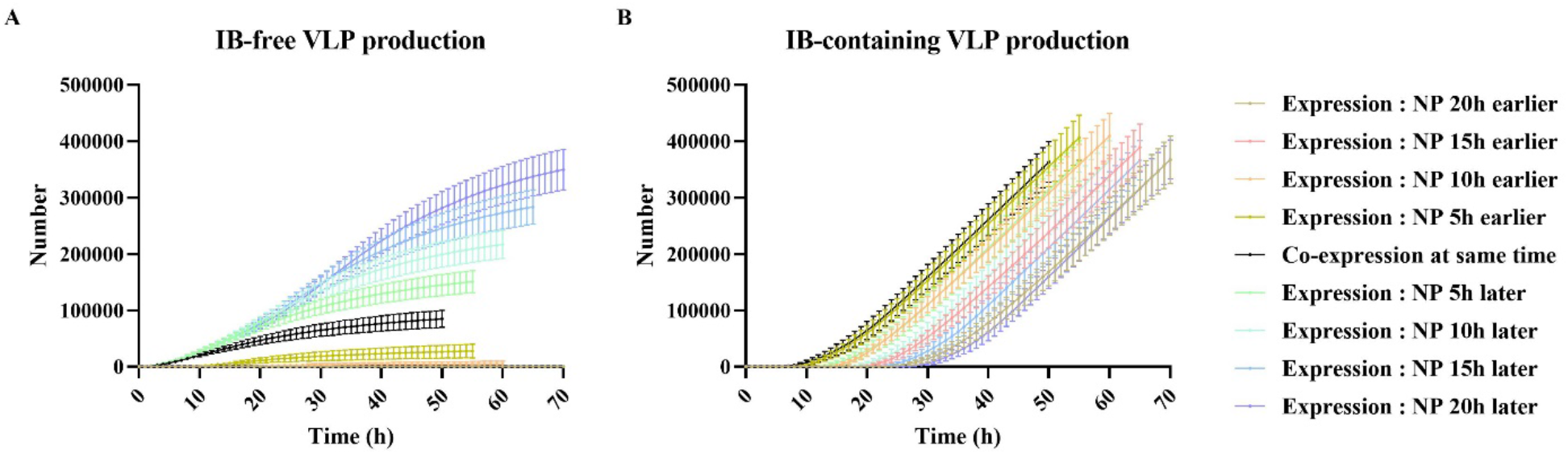
Influence of NP/VP40 expression time on VLP production. (A) IB-free VLP production decreases as VP40 expression time becomes later. (B) IB-containing VLP production decreases expression time difference between NP and VP40 becomes larger. The highest IB-containing VLP production appears when VP40 expression time is between 0-5h later than NP expression time. Error bars indicate 95% CI. 41 out of 50 groups are used for analysis as others have met tolerance problems in ode-solver.

**Figure 8.**
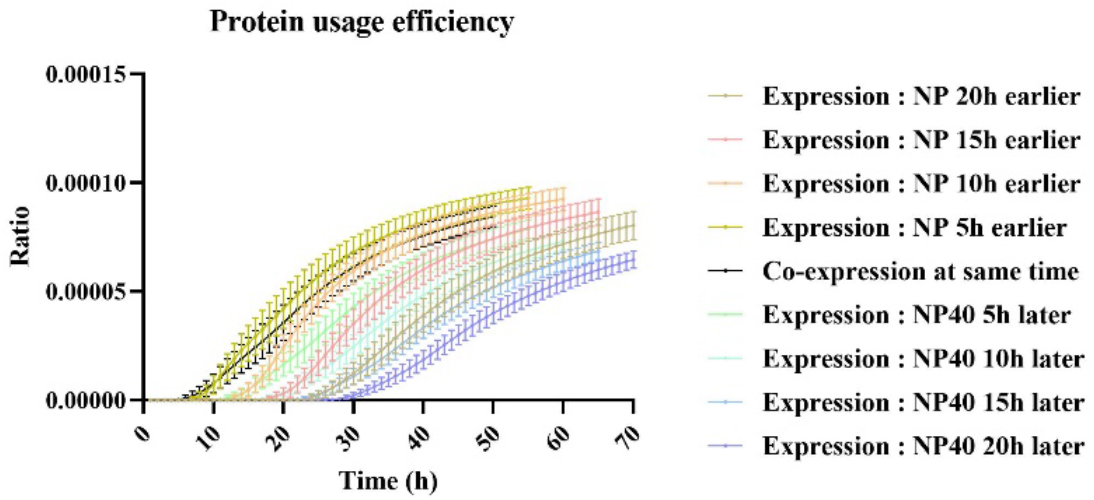
Ratio of IB-containing VLP production to protein production. The ratio of IB-containing VLP to total protein production is a representative of energy use. It reaches the peak when VP40 is 5 h late in expression, and either earlier or later expression time will decrease the energy usage. Error bars indicate 95% CI.

In conclusion, both the expression ratio and the expression time of NP and VP40 are important for the effective production of IB-containing viral particles. An optimum ratio and expression time exists for producing viral particles in the shortest time and for reducing energy waste by suppressing noninfectious viral particles. From our study, we believe if the protein expression profile is disturbed in EBOV infection, such as having VP40 expressed earlier than NP, the production of infectious viral particles can be significantly impaired.

### Inhibition of fendiline on VLP production is weakened when NP is co-expressed

Finally, as we had evaluated fendiline as a potential small-molecule drug for EBOV treatment by VP40 computational modeling in our recent work, we wanted to see how fendiline treatment influenced the NP-VP40 system. We simulated fendiline treatment by changing its concentration input (see Methods) Our results indicate that both IB-free and IB-containing VLP production is decreases with fendiline concentration (Fig. S4, Table S15). This indicates that fendiline can be effective at suppressing VLP production when NP is co-expressed with VP40. However, if we compare the reduction percentage in VLP between NP co-expression and VP40-only, we find that the inhibition of VLP production by fendiline is weakened in the NP-VP40 system (Fig. 9, Table S16). Considering our recent results (25) that fendiline is less effective when the VLP budding rate is high, we believe this is caused by the higher budding rate of IB-containing VLPs. Since VLP production can be improved by multiple EBOV proteins (13), fendiline treatment efficiency may be lower in authentic EBOV infection than our predictions from simulations. This is also aligned with experimental findings that fendiline is less effective against live EBOV than VP40 VLP under the same fendiline concentration (28). A co-treatment targeting the budding process of EBOV may be important to rescue the efficiency of fendiline, as suggested in our recent work (25).

**Figure 9.**
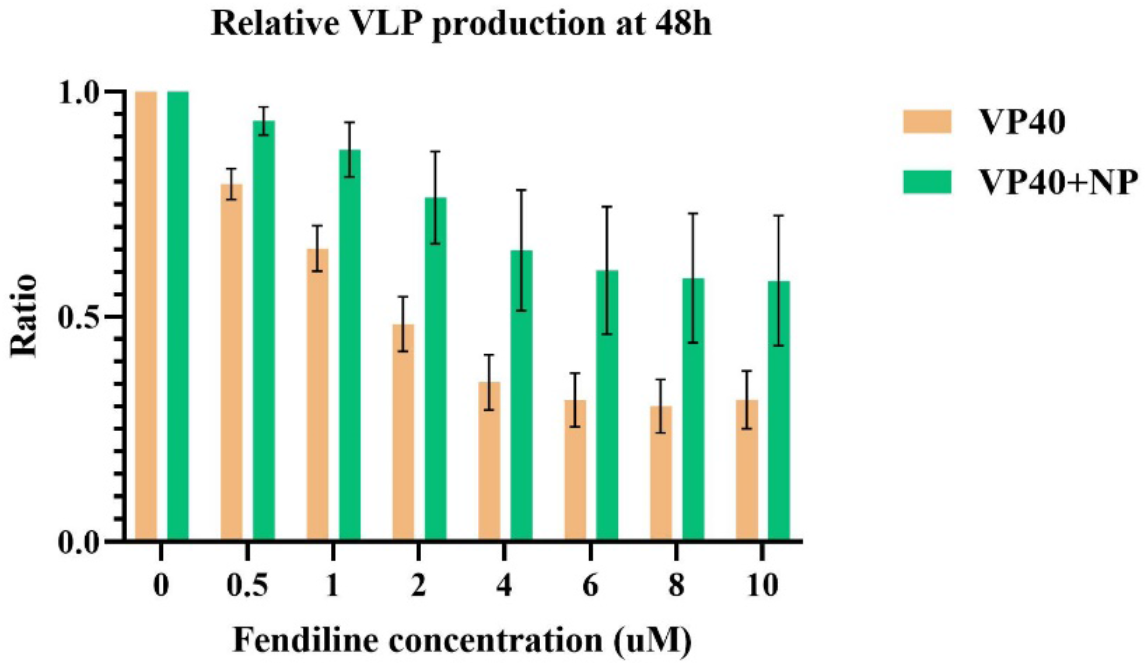
Inhibition of VLP production by fendiline on VP40-only and NP-VP40 system. While total VLP production is inhibited in both VP40-only and NP-VP40 system by fendiline, the reduction in VLP is smaller in NP-VP40 system. Error bars indicate SD.

## Discussion

NP is an important viral protein for the whole life cycle of EBOV. In this study, we have incorporated NP into our ODE-based VP40 assembly and budding model by a two-stage interaction mechanism and explored the detailed impact of NP on VLP production through computational methods.

A recent study found that the interaction between VP40 and NP can happen in both cytoplasm and membrane (17). We believe that the IB-bound VP40s and cell membrane-bound VP40s are two different pools. The former will change the structure of NP-CTD by interacting with NP-NTD in the cytoplasm, and the latter will move to cell membrane, where it recruits IBs and serves as building block for filaments. We have successfully replicated experimental observations through our model based on these mechanisms, which also supports the two-stage interaction theory between NP and VP40. Though a previous study indicated no interaction between VP40 and NP in the cytoplasm (22), the study was conducted in Marburg virus (MARV) instead of EBOV, and NC was used instead of NP alone as in the current study. The difference in species and the discrepancy between NC and IB may explain the observed differences in NP-VP40 interaction. We can further explore a system without cytoplasmic NP-VP40 interaction in our model to assess potentially different mechanisms between different filoviruses.

While a previous study found that VLP production can be increased by co-expression of NP and VP40 (13), our simulations further suggest that the enhancement may be due to the stabilization of growing VP40 filaments and increase in VLP budding rate through IB association. These influences will increase VLP budding in general, while reducing the production of IB-free VLPs, which is aligned with experimental observations (13, 17). However, our results also indicate that too many NPs may inhibit VLP production by depleting membrane bound VP40 dimer building blocks in two ways: (a) Too many cytoplasmic NPs will bind more VP40s and trap them in cytoplasm; (b) The stabilized IB-containing VP40 filaments are less likely to release VP40 dimers. Taken together, it means apart from having mutants in the NP CTD (17), NP may also serve as a “trap” for VP40 if the number of NPs is too large since part of VP40s will be limited to NP NTDs, and the rest of VP40s will be distributed between a large number of incomplete VLPs that are not mature enough to bud. Since we lack data on how many VP40s will interact with NPs in the cytoplasm, we aren’t able to determine a precise ratio of NP/VP40 where it will start inhibiting VLP production. This could be further quantified through experiments.

On the mRNA level, NP shows a similar level of transcription compared to VP40 (29, 30). On the other hand, the size of NP (739 aa) is more than twice that of VP40 (326 aa), so it can be inferred that VP40 translation time should be shorter while not considering the difference in amino acid elongation rate caused by codons. Thus, VP40 may be more abundant than NP in EBOV infected cells. Since NP is at the 3’ of EBOV genome, the expression should be earlier than VP40. mRNA detection also shows that the level of NP transcription decreases through time (29). Our simulation indicates that these experimentally observed expression patterns of NP/VP40 timing and ratio are beneficial to both the production of EBOV particles and the suppression of non-functional EBOV without genetic material. It maximizes IB-containing VLP production, while decreasing energy waste. These results suggest that the expression profile of individual EBOV proteins may play a critical role in its life cycle, and thus can be a potential treatment target for future studies. Currently, no RNA-based therapy has been approved by the FDA in treating EBOV infection, as all approved treatments of EBOV are antibody-based (5, 6). However, RNA interference (RNAi) has been proposed for viral infection treatment for many years and is considered an efficient means of disruption (31). Also, there are already experimental therapy using small interfering RNA (siRNA) therapy targeting multiple EBOV proteins (32, 33), though the treatment effect remains challenging (34). The difficulty lies in accurate delivery to target cells (31, 34, 35), and the efficiency may be affected by application time (34). However, RNA delivery technology has been greatly advanced recently (36, 37), characterized by the recent mRNA vaccines approved for COVID (38, 39). Our previous work also showed the ability of computational models to assist evaluation of treatment timing (25). Taken together, our results further suggest that an RNA-based therapy which can disrupt the normal relative expression abundance of EBOV proteins over time could significantly impact infectious virus production.

Our results highlight the power of simplified *in vitro* and *in silico* models to disentangle complex protein-protein interaction network structures and dynamics. However, from the fendiline results in this study, we conclude that the interaction between various EBOV proteins can influence treatment efficiency. Thus, our findings also caution against extrapolating drug target conclusions made in simplified *in vitro* or *in silico* systems that only consider one viral protein. It is important to build the full EBOV infection model for making more accurate efficiency predictions. Our current model still has limitations. The current model does not include viral entry, transcription, replication or the five other EBOV proteins. For the NP-VP40 model, we lack the knowledge of what part of NPs can interact with and with how many VP40s on IBs in cytoplasm, and thus our model represents a simplified assumption. We also have limited data on parameter values, or enough data to calibrate them precisely. Nonetheless, our NP-VP40 model successfully reproduces current experimental results, makes important predictions, and provides valuable directions for future experiments.

In summary, our results show how EBOV NP affects viral assembly by influencing filament stability and buddy rate constants; how the timing and proportion of NP vs. VP40 expression from the viral genome is potentially optimized for maximal functional virion production; and how viral protein interactions impact drug efficacy. Thus, this work moves the field forward in our understanding of EBOV assembly dynamics; and brings us one more step towards a full EBOV infection model, which can be used for *in silico* treatment trials.

## Materials and Methods

### ODE-based model construction

Our model incorporates NP dynamics into our existing model of VP40 assembly and budding (r_1_,r_2_, k_1_, k_2_, k_3_, k_4_, k_5_, d_1_, d_2_) (24, 25). NP is produced and assembled into IBs in the cell cytoplasm (r_1_, k_6_, d_1_), and then bound to VP40 dimer at the plasma membrane (k_8_) (as experimental evidence has not shown whether IB binds to VP40 membrane dimer or higher oligomers, we make a simplifying assumption that excludes the interaction between IB and higher VP40 oligomer). Cytoplasmic NPs also have the ability to incorporate cytoplasmic VP40 dimers (k_7_). The ability of IBs to bind to membrane VP40 depends on the cytoplasmic VP40s attached to it (k_8_) (17). Finally, membrane VP40 can oligomerize into either IB-free filaments or IB-containing filaments (k_3_, k_4_, k_9_, k_10_) (Fig. 10).

**Figure 10.**
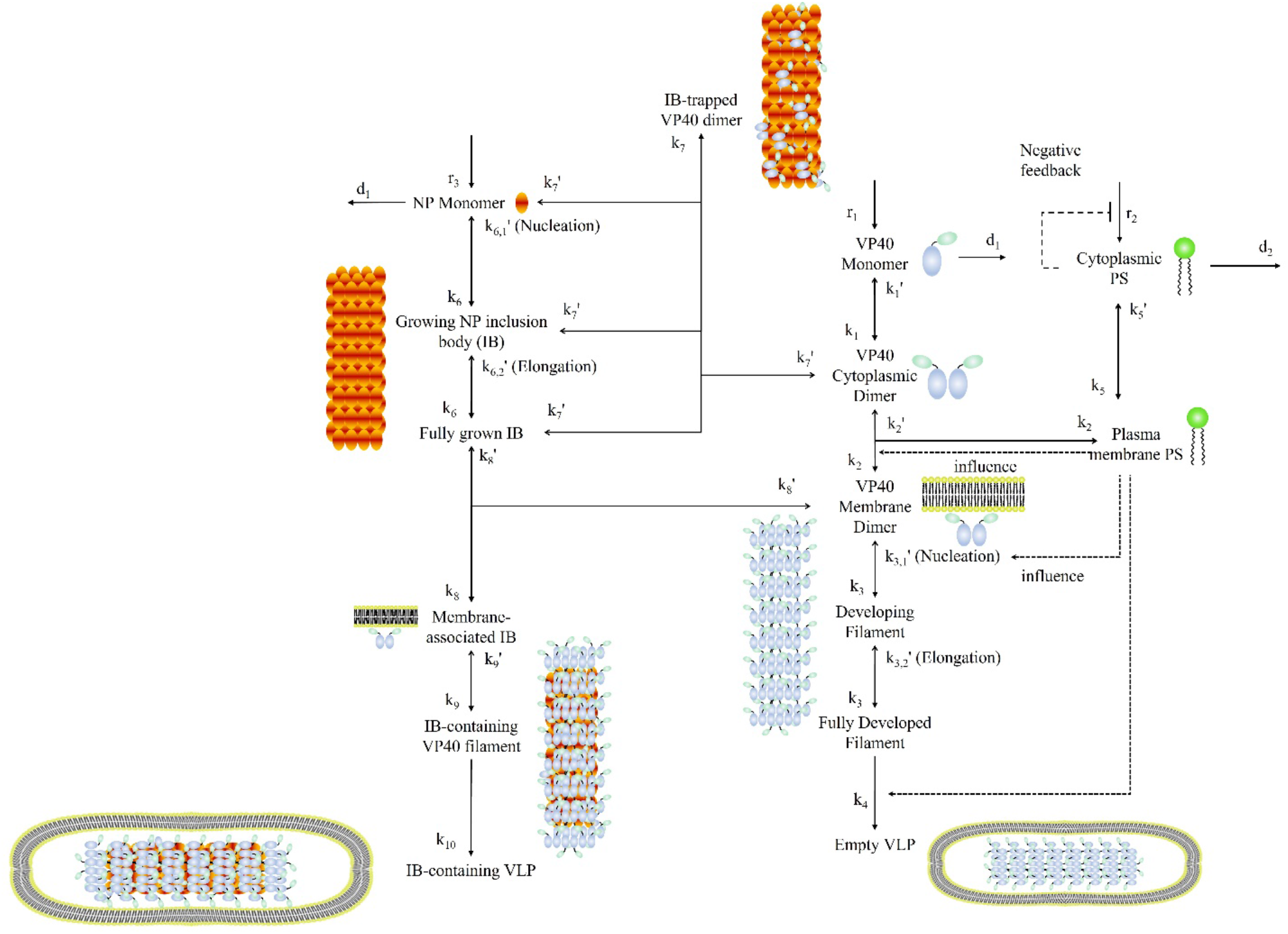
Scheme of the NP-VP40 model. The NP-VP40 model includes the pathway from NP production to IB formation, VP40 assembly and budding (both IB-free and IB-containing) as well as simple PS metabolism network.

ODEs for the model are shown in Eq. (1)-(24). Eq.(1), Eq.(2), Eq.(8) and Eq.(10)-(15) are being reproduced here from our prior work (25).

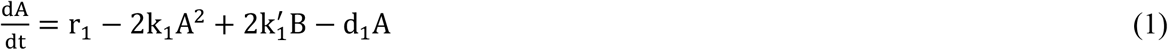

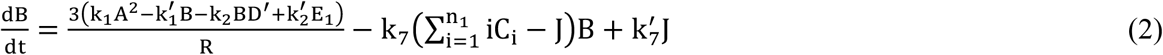

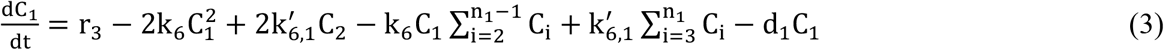

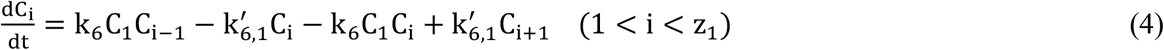

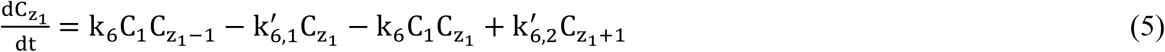

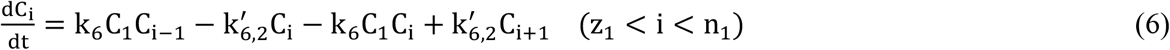

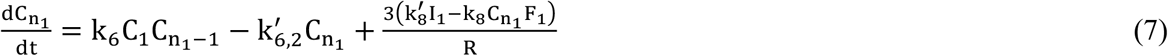

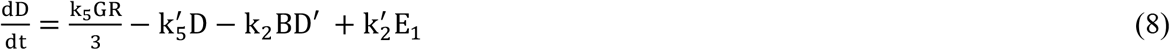

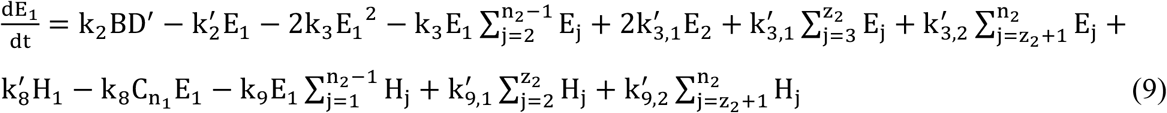

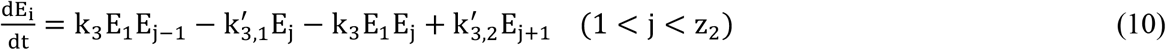

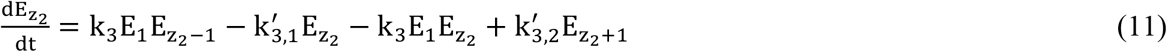

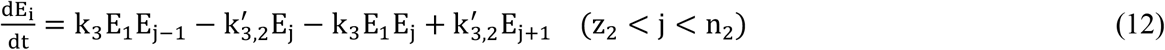

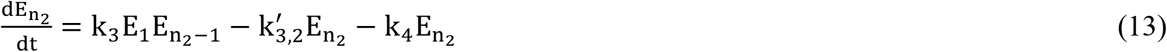

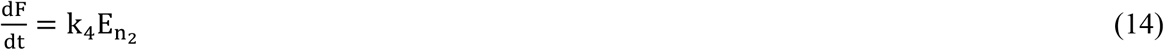

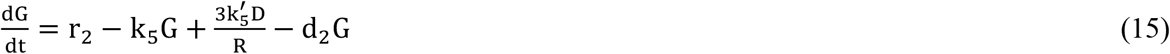

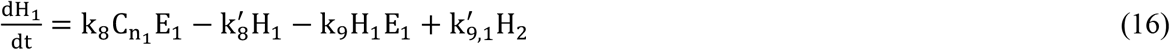

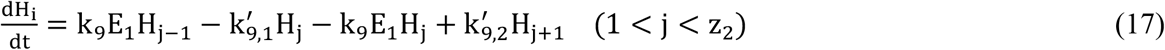

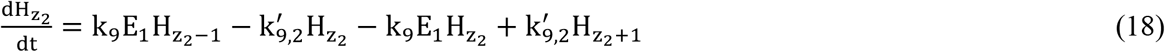

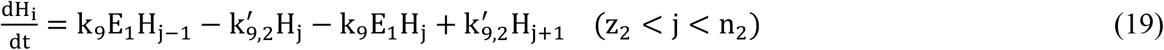

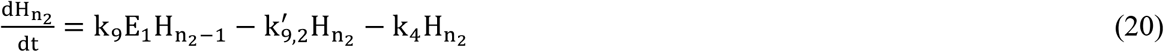

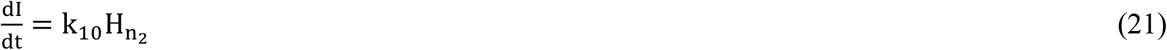

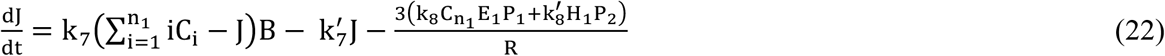

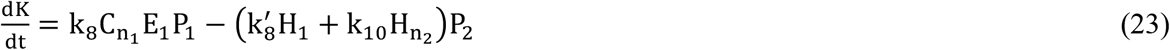

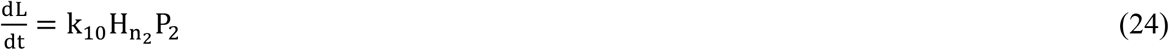

A: VP40 monomer in cytoplasm (nM). B: VP40 dimer in cytoplasm (nM).

C_i_: Developing cytoplasm IB consists of i NPs (nM). i: Number of NPs in cytoplasm IB.

z_1_: Size of IB where the reverse rate constant change from k_6,1_′to k_6,2_′ (from nucleation to elongation

n_1_: Number of NPs contained in a mature IB. n_1_ = 770 in our model. D: Plasma membrane phosphatidylserine (nmol/dm^2^).

D′: Plasma membrane Phosphatidylserine available to interact with cytoplasmic VP40 dimer (nmol/dm^2^).

E_j_: Developing IB-free VP40 filament consists of j VP40 dimers (nmol/dm^2^). j: Number of dimers in developing filament.

z_2_: Size of VP40 filament where the reverse rate constant changes from k_3,1_′to k_3,2_′ (from nucleation to elongation).

n_2_: Number of dimers in a mature filament. n_2_ = 2310 in our model. F: Budded IB-free VLP (nmol/dm^2^).

G: Cytoplasmic phosphatidylserine (nM).

H_j_: Developing IB-containing VP40 filament consists of j VP40 dimers (nmol/dm^2^). I: Budded IB-containing VLP (nmol/dm^2^).

J: Cytoplasmic VP40 dimer trapped in cytoplasm IB (nM). K: Trapped VP40 dimer in plasma membrane IB (nmol/dm^2^).

L: Trapped VP40 dimer in budded IB-containing VLP (nmol/dm^2^). P_1_: Portion of cytoplasmic NP bound by VP40 dimer.

P_2_: Portion of plasma membrane NP bound by VP40 dimer. P_3_: Portion of budded NP bound by VP40 dimer.

Definitions of P_1_-P_3_ are shown in Eq. (25)-(27).

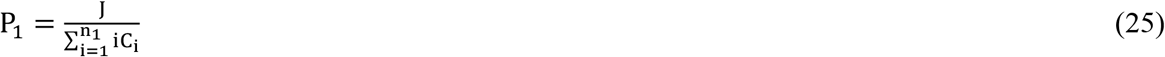

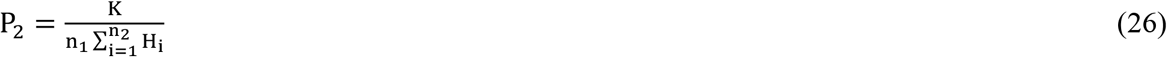

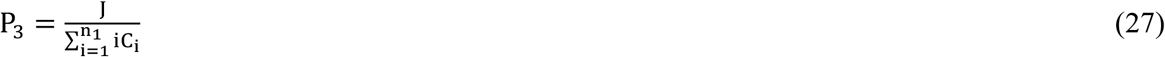

Initial conditions:

A(0)=0

B(0) = 0

C_i_(0) = 0

D(0) = 16.75

D_i_(0) = 0 (1 ≤ i ≤ n_1_)

E_j_(0) = 0 (1 ≤ j ≤ n_2_)

F(0) = 0

G(0) = 1.07 × 10^5^

H_j_(0) = 0 (1 ≤ j ≤ n_2_)

I(0) = 0

J(0) = 0

K(0) = 0

L(0) = 0

The calculation of D’ is shown in Eq. (28). The deduction of the equation is detailed in our prior work (25).

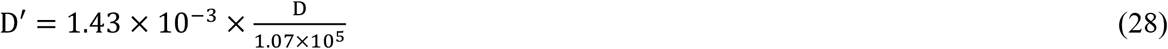

### Influence of IB-bound VP40 on IB cell membrane association

Previous work has shown that the interaction of NP NTD with cytoplasmic VP40 can cause a conformational change in the NP CTD, which is critical for the recruitment of IB to cell membrane (17). We reflect this mechanism in our model, by having the IB membrane association rate (k_8_) positively impacted by the portion of its NP occupied by cytoplasmic VP40 dimer. The influence is described in Eq. (29).

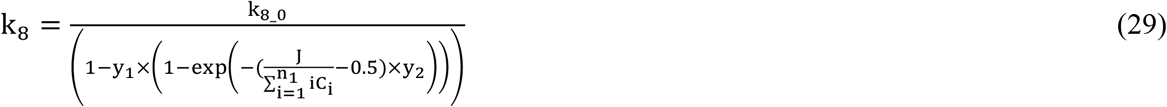

Values of y_1_ and y_2_ are listed in Table 1.

**Table 1.**
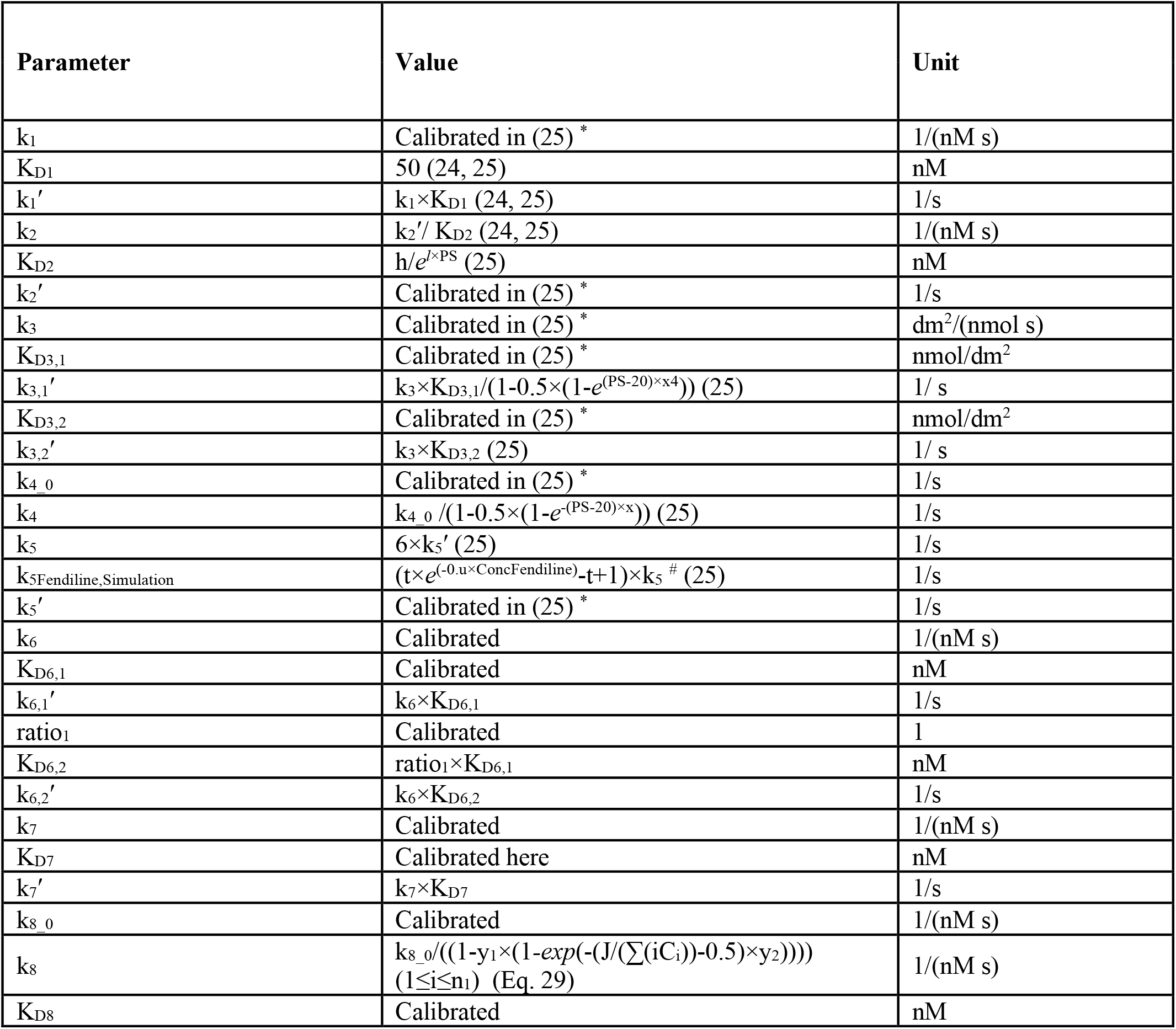
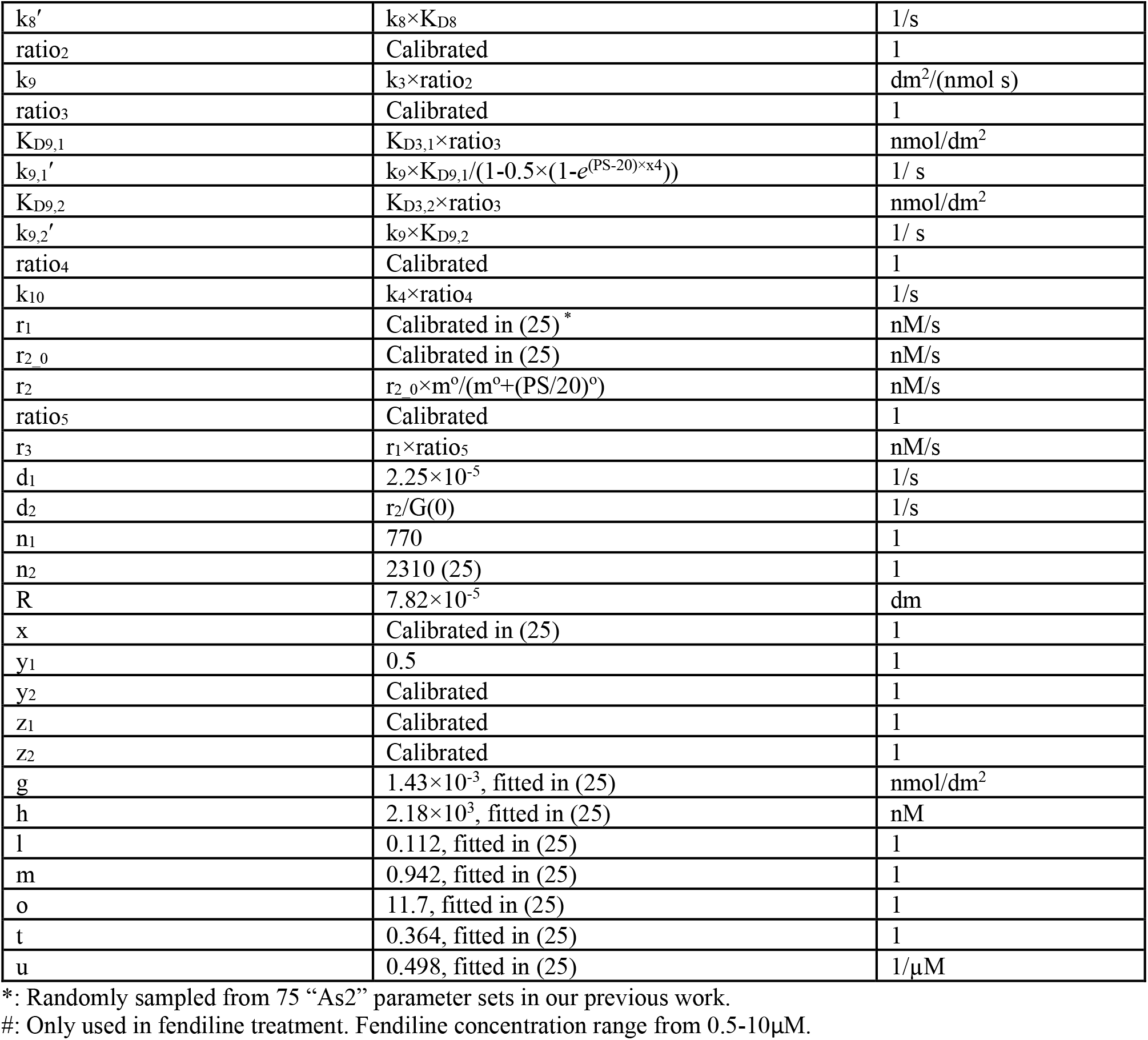
Model parameters.

k_8_0_: Calibrated IB plasma membrane association rate without considering the effect of attached cytoplasmic VP40 (Table 1).

### Experimental data

Three groups of data from two NP-VP40 experimental studies are used to calibrate our model:

- NP VLP production ratio: defined as the ratio of VLP production with NP co-expression relative to VLP production with expression of VP40 alone (13).
- CTD-mutant NP (L692A, P697A, P698A, W699A, which have CTD mutations and are compromised in binding to the cell membrane) VLP production ratio: defined as the ratio of VLP production with wild-type (WT) NP co-expression with VP40 relative to CTD-mutant NP co-expression with VP40 (17).
- CTD-mutant NP membrane VP40 ratio: defined as the ratio of cell membrane VP40 number relative to cytoplasmic VP40 number in both WT and CTD-mutant NP co-expression (17).

CTD-mutant NP data are combined together in our calibration since they are all mutated in the CTD core and are also analyzed collectively in the experimental work (17). We reflect this mutation mathematically by letting k_8_ = 0 for these mutants. All the data are included in Table 2.

**Table 2.** Calibration data. SD: standard deviation.

|  | NP VLP production ratio | Mutant NP VLP production ratio | Membrane VP40 ratio |
| --- | --- | --- | --- |
| WT NP+VP40 | 3.6078 (SD: 1.961) |  | 6.325 (SD: 1.554) |
| Mutant NP+VP40 |  | 0.3145 (SD: 0.1274) | 0.637 (SD: 0.142) |

As qualitative validation of the NP-VP40 model, we evaluate the fitted parameters by their ability to reproduce the observed dynamics that when NP is expressed by itself, the average IB size increases over time; and that the IBs have a bimodal size distribution, with the majority of IBs being either very large or very small at latter time (26).

### Calibration and parameter estimation

NP-related parameters are sampled through Latin hypercube sampling (LHS) in a wide range, while VP40-only parameters are randomly sampled from the 75 “As2” simulation groups in our last study (25). The agreement of model predictions with experimental data is calculated by a cost function as described in our last study and shown in Eq. (30). Compared to calculations in our previous work, we removed the weight that was introduced to account for the uncertainty in some data sets. All predictions at a certain time point are calculated from the average value within ± 2 hours as in our previous work (25).

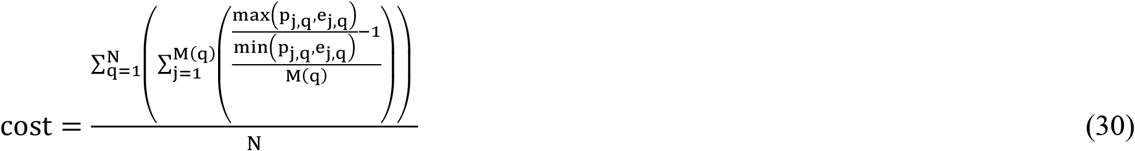

N: Number of different data type

M(q): Number of data in the qth data type

e_j,q_: jth experiment data in the qth data type

p_j,q_: jth model prediction in the qth data type

Calibration is performed in an iterative manner. In each iteration, 2500 initial guesses are sampled with LHS. The top 50 parameter sets with the lowest costs are used for determination of the parameter ranges of next iteration. Since experimental studies showed that IB-containing VLPs are dominant compared to IB-free VLPs at 36h post-transfection (17), we also use this feature to filter parameter sets for further analysis starting from the 3^rd^ iteration (such cases are very limited in iterations 1-2, Table S1). The ranges of calibrated parameters in each round are shown in Table S2. Calibration is considered complete after six iterations due to an increase in cost in the seventh iteration compared to the sixth. Some parameter sets have P_1_-P_3_ higher than 1 at some time points due to numerical errors and are excluded from further analysis and simulation. The top 50 best fit parameter sets from the rest of the sixth iteration samples are selected for further analysis (Table S3).

### Simulation

Three kinds of simulations are performed: local sensitivity analysis, expression time test and fendiline treatment. In local sensitivity analysis, selected parameters are changed from 0.1× - 10× of their original values, while other parameters remain the same. These local sensitivity analysis allows us to evaluate how the impact of NP on different steps in VP40 assembly and budding process can influence VLP production, as well as the importance of the NP/VP40 expression ratio. In expression time tests, the relative monomer production starting time of NP ranges from 20h earlier than VP40 to 20h later. The expression time tests will enable us to assess if the relative expression time of NP/VP40 is important in EBOV infection. In fendiline treatment, 0.5 µM - 10 µM of fendiline is simulated in the NP-VP40 system to test how disruption of phosphatidylserine by fendiline affects VLPs when NP is co-expressed with VP40.

### Software application

The ODE model is implemented in Matlab R2022a. Solver “ode23t” is used for solving ODEs, with the analytical Jacobian Matrix provided, and using the “NonNegative” setting to avoid negative values. All result figures and statistical analysis are created with Graphpad Prism.

## Funding acknowledgements

This project was funded with support from the Indiana Clinical and Translational Sciences Institute which is funded in part by Award Number UM1TR004402 from the National Institutes of Health, National Center for Advancing Translational Sciences, Clinical and Translational Sciences Award (to EP and RVS); and AI081077 (to RVS). The content is solely the responsibility of the authors and does not necessarily represent the official views of the National Institutes of Health. This material is based upon work supported by the National Science Foundation under Grant No. 2143866. This work used the Extreme Science and Engineering Discovery Environment (XSEDE), which is supported by National Science Foundation grant number ACI-1548562. Anvil at Purdue was used through allocation TG-MDE220002 (to EP).

